# Integrated whole-genome and transcriptome sequencing reveals divergent evolutionary processes across biliary tract cancer subtypes

**DOI:** 10.64898/2025.12.12.693962

**Authors:** Felix E.G. Beaudry, Duhan Yendi, Danielle Arshinoff, Nicholas Light, Simona Perrotti, Erin Winter, Liam R. Cristant, Jingxiong Xu, Julie Wilson, Anna Dodd, Roxana Bucur, Eric X. Chen, Elena Elimova, Rebecca Wong, Aruz Mesci, Ali Hosni, Anand Ghanekar, Raymond Jang, Chaya G. Shwaartz, Trevor Reichman, Carol-Anne Moulton, Enrique Sanz-Garcia, Grainne M. O’Kane, Erica S. Tsang, Xin Wang, Ian McGilvray, Steven Gallinger, Trevor J. Pugh, Gonzalo Sapisochin, Arndt Vogel, Jennifer J. Knox, Faiyaz Notta, Robert C. Grant

**Author notes:** Correspondence: Robert C. Grant.

## Abstract

**Introduction:** Biliary tract cancer (BTC) comprises a family of rare malignancies subclassified by anatomy and pathology. However, this scheme may obscure shared biology and limit patient stratification.

**Objectives:** We tested whether BTC heterogeneity can be explained by coherent latent axes and evaluated the potential to unify diverse clinical and genomic factors under a tractable biological framework

**Methods:** We performed whole-genome and transcriptome sequencing of 180 tumors enriched for tumor cells by laser capture microdissection to identify shared programs in BTC.

**Results:** Network integration across transcriptomic classes identified two consensus cancer subtypes (CCS). CCS segregated with anatomical location of primary tumor and gene expression marker analyses suggest subtypes reflect tumor cell of origin differences. CCS displayed strikingly divergent molecular landscapes, explaining more variance than anatomical location of primary tumor. CCS-B tumors were mutationally loaded with clock-like and APOBEC signatures and extrachromosomal DNA, whereas CCS-A tumors were characterized by chromosome-arm deletions.

**Conclusion:** Our approach showed that harnessing the genomic and transcriptomic diversity of BTC uncovers novel biology and improves stratification.

**Significance:** We provide evidence that biliary tract consensus cancer subtypes define fundamentally different cancers, with diverging modes of evolution stemming from distinct cells of origin.

## Introduction

Biliary tract cancer (BTC) is a rare and highly fatal disease ^1^. Progress in understanding treatment response and identifying actionable biomarkers has been hindered by its extensive clinical and molecular heterogeneity ^2,3^. The different staging across BTCs showcases its heterogeneity, with distinct systems for intrahepatic, perihilar, distal, and gallbladder cancers. The multitude of anatomical, molecular, and etiological factors makes it difficult to isolate the impact of any single variable, limiting the interpretability of traditional comparative analyses.

Emerging evidence suggests that many of these heterogeneous features covary, raising the possibility that they reflect deeper, latent organizational principles within BTC biology. Certain driver mutations tend to co-occur more than others, such as *FGFR2* fusions with *BAP1* mutations or *KRAS* with *TP53* mutations ^4^. The former tend towards intrahepatic tumors while the latter appear most in extrahepatic tumors. Patients with primary sclerosing cholangitis (PSC) or liver fluke infections, both of which are leading BTC risk factors, tend to have *KRAS* or *TP53* mutant tumors ^5,6^. Several gene expression classifiers with prognostic implications show different rates of driver mutations ^7–11^, but how they relate to each other remains underexplored.

To date, clinical BTC classification has mostly been anchored to anatomic origin or mutations in specific genes ^12^. In this study, we tested whether BTC heterogeneity can be explained by coherent latent axes and evaluated the potential to unify diverse clinical and genomic factors under a tractable biological framework.

## materials and methods

### Patients and cohort

Patients included in this study contributed to the LeGresley Biliary Registry. This registry prospectively recruits consentable adults over 18 years of age, diagnosed with BTCs at the University Health Network, including individuals with advanced-stage disease. Eligible patients were identified for recruitment through a systematic review of clinic lists. Demographic and clinical data were prospectively collected from the time of enrollment. TNM staging was performed according to AJCC 8^th^ edition. Sample and basic data processing methods can be found in the supplementary materials.

### Ethics approval and consent to participate

Ethics approval was granted by the University Health Network institutional review board (REB# 24-5412, approved September 11, 2025), and informed consent was obtained from all subjects in this study. All processes were conducted in accordance with the Declaration of Helsinki.

### Expression class network and consensus subtyping

For all RNA samples, transcripts were summed at the gene level using annotations from Ensembl release 100. Count data were normalized using *Deseq2* v 1.48.1 ^41^ and scaled using *ematAdjust* from *CMScaller* v0.99.2 ^42^. Samples were assigned subtypes using Nearest Template Prediction (*ntp*) with *CMScaller* over 1,000 permutations. Where the false discovery rate of class assignment of a sample was greater than 0.05, the class assignment for that classifier was considered undefined.

To build a consensus cancer subtype (CCS) network, we included 23 signatures from nine previously published BTC tumor classifiers (Table S1). Signature similarity was assessed using a Jaccard similarity matrix. Classifiers (nodes) were removed where (1) a classifier had a lack of conclusive assignment in >50% of cases or (2) was connected to the rest of the network by less than 2% of samples. Additional signatures were removed in optimization of the network’s inflation factor maximizing modularity and silhouette scores, evaluating values between 1.5 and 20 by steps of 0.5 (Fig. S2c). Network clusters were identified with the Markov cluster algorithm *mcl* v 1.0 ^14^.

To train a top-scoring pair algorithm, we restricted the training set to samples that were assigned to the same CCS across every classifier. Classifiers where only some classes were included in the CCS, and therefore led to less than 50% of samples assigned to a CCS, were excluded from TSP training. The algorithm was trained on 80% of the data (*n* = 70) with the *SWAP.KTSP.Train* function of *switchBox* v 1.44.0 ^15^ in the range from 10 to 16 genes pairs and tested on the remaining 20%.

To train a copy number-based CCS classifier, the top 10 arms loading onto the first principal component of inter-sample arm-level relative copy number alteration variance were included in a generalized linear model and arms were dropped stepwise to minimize the number of factors in the model using the *step* function from *stats* v 4.5.1. Given the use of relative copy number, incorporating the mean variant allele frequency of exonic mutations (as a proxy for tumor cellularity) significantly improved the fit of the model. For external validation of the gCCS classifier, data from Farshidfar *et al.* ^16^ were downloaded from cbioportal.org. Methods for processing of external datasets can be found in supplementary materials.

### Gene expression profiles and virtual microdissection

Differential gene expression between samples from each substype was evaluated using *DESeq2*. Non-negative matrix factorization (NMF) was performed 200 times (nrun = 200) on RNA read counts per million minimizing the Frobenius norm using *NMF* v 0.28. To identify malignant components, we normalized component weights, subset to samples where weight was larger than zero for that component and filtered for components that had a significant correlation with genomics-estimated tumor content greater than 0.2. Optimization of k was performed by finding k where both the average correlation between signature weight and tumor content across components was maximized. NMF weights were converted to pseudo transcript per million values for comparison among samples. Estimates of tumor content from pathologists was performed on 39 samples, and correlation was only estimated on cases for which LCM was not performed as pathologists did not re-evaluate tumor content between LCM and sequencing.

Single cell RNAseq biliary duct cell type markers were downloaded from Table S1 from Sampaziotis *et al.* ^22^ and single nucleus data from Table S10 from Andrews *et al.* ^43^ and run on *GSEA* from *clusterProfiler* v 4.16.0 ^44^. Human Protein Atlas enrichment was performed using *gost* from *gprofiler2* v 0.2.3. *CIBERSORT* correlations were obtained from *immunedeconv* v 2.1.0 ^45^ deconvolution with “cibersort_abs”.

### Genomic variant calling

We used *ActiveDriverWGS* v 1.2.1 ^46^ to look at statistically recurrent small mutations while controlling for variation in mutation rate across the genome. To look more directly for signals of selection in protein-coding genes, we also used *dNdScv* v 0.0.1.0 ^47^. Promoter regions were tested exclusively with *ActiveDriverWGS*. All significantly recurrent hits were manually reviewed to exclude germline variant contamination.

Expression quantitative trait loci were identified using a linear regression model for transcript per million as a function of the presence of a structural variant within 5 kb of the gene of interest, using p-values adjusted with Benjamini-Hochberg correction. Breakpoints were tested for statistical recurrence using *FishHook* v 0.1 ^33^. Driver co-occurrence significance was corrected for variant frequency using *Rediscover* v 0.3.2 ^48^.

Fisher’s exact test queried whether genes were enriched for loss-of-heterozygosity when hit by a substitution. *GISTIC* v2.0 ^34^ was used for testing recurrent copy number alterations and converted into gene-level scores using *CNtools* v 1.64.0. To isolate the functional consequences of copy number alterations, we filtered genes in recurrently altered regions for a significant correlation between relative copy number and gene expression measured in transcripts per million.

### Mutational landscape

Mutational signature analysis was performed *de novo* using SigProfilerExtractor v 1.2.2 ^49^, with standard parameters, and decomposition into COSMIC v3.4 reference signatures performed using non-negative least squares. Signature analysis was performed separately for SBS, DBS and In/Dels. eCCS signature enrichment was performed using a binomial linear regression model ^50^, with the response as a binary variable representing the appearance of a base assigned to that signature in the given sample, filtering for signatures present in more than 30% of samples.

### Code availability

Full code walkthrough as well as single-sample CCS classifiers are found at pancurx-oicr.github.io/btc-ccs/.

### Data availability

All raw WGS and WTS newly generated for this paper are available on EGA under dataset ID EGAS50000000972. Sample metadata and processed matrices can be found in supplementary materials (Table S7-9).

## Results

### Cohort description

We analyzed 180 cases of BTC, including 111 new cases recruited prospectively and 69 previously reported cases ^12,13^. The median age of patients was 61 years (range 29-83) and tended towards males (*n =* 107). 97 patients underwent curative-intent resections, followed by adjuvant capecitabine (*n =* 43), and 122 received palliative-intent systemic therapy, most commonly gemcitabine, cisplatin, and durvalumab (*n =* 74; Fig. 1a, S1a); clinical course information was unavailable for 11 patients.

**Fig. 1.**
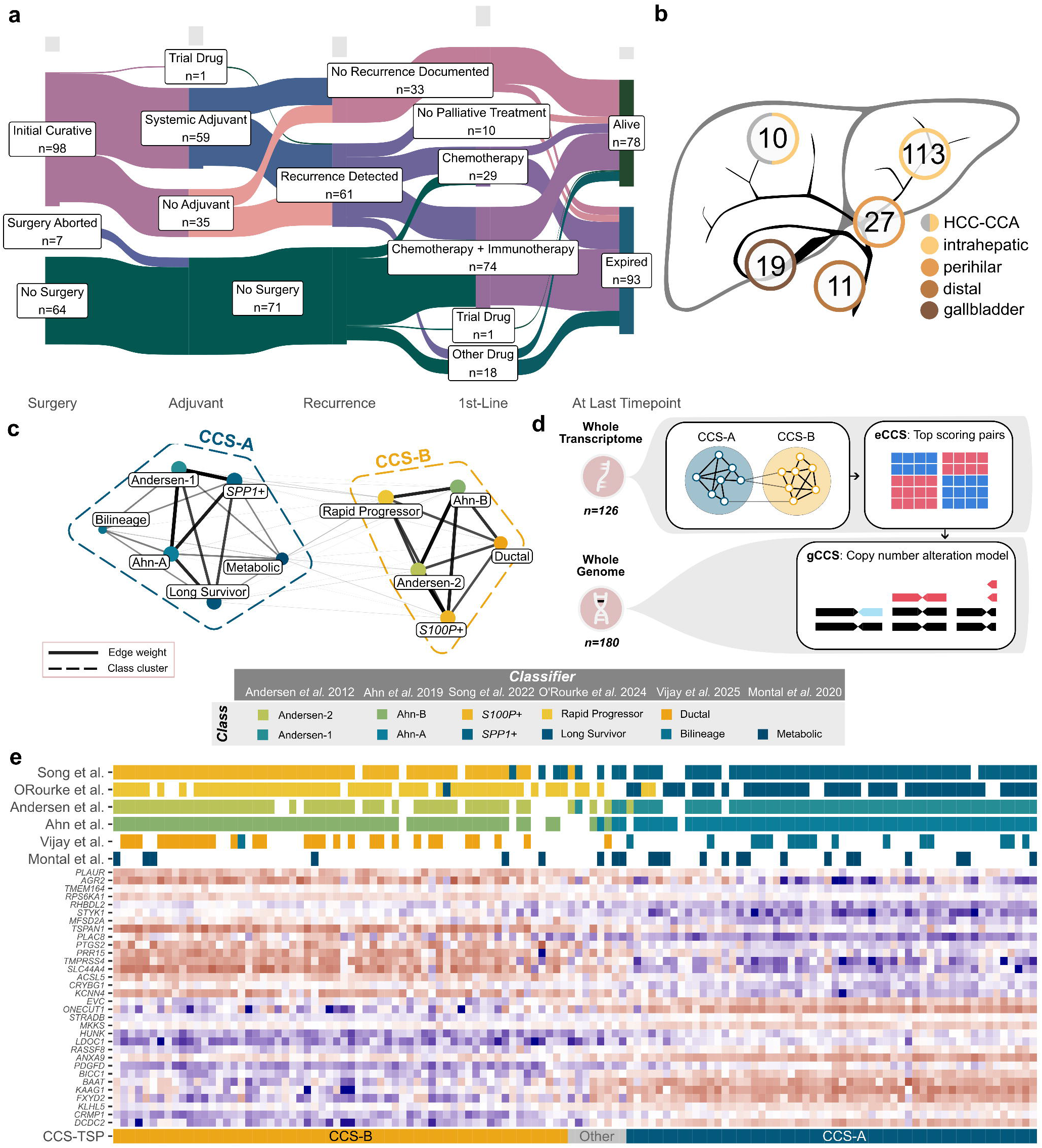
Defining consensus cancer subtypes of biliary tract cancer. **a**, counts of patients proceeding through different clinical trajectories. **b**, counts of primary anatomical tumor site. **c**, assignment similarity network between the CCS. **d**, classifier method schema. **e,** Assignment of fresh frozen whole transcriptome samples to 6 published classifiers, as well as a heatmap of the 16 genes in the classifier and the derived CCS-based top-scoring pair classifier. HCC-CCA, combined hepatocellular-cholangiocarcinoma. CCS, consensus cancer subtype. eCCS, expression-based consensus cancer subtypes. gCCS, genetics-based consensus cancer subtypes. TSP, top scoring pairs.

The whole genome (WGS) of one tumor specimen was sequenced for every case and underwent whole transcriptome sequencing (WTS) when enough tissue remained (*n* = 145; Fig. S1b). As BTCs typically exhibit desmoplasia and low tumor cellularity, most samples were enriched for tumor using laser-capture microdissection (*n* = 164). Fresh frozen samples (*n* = 159) were preferred over formalin-fixed paraffin-embedded tissue (*n* = 21). Primary tumors spanned the biliary tree, including 113 intrahepatic, 10 combined hepatocellular-cholangiocarcinoma, 27 perihilar, 11 distal, and 19 gallbladder tumors (Fig. 1b). Metastases were detected in 66 patients (40% of evaluable); 32 specimens were from metastases, including 24 liver biopsies from extrahepatic cancers (Fig. S1c, d). Half of tumor specimens (n = 70, 51%) were conclusively treatment-naive (Fig. S1b).

### Network integration of published transcriptomic classifiers reveals two subtypes

Expression-based classifiers of BTC have been previously proposed but studied in isolation. To explore overlap between previous classifiers, we classified all 126 fresh frozen samples with WTS into 23 classes from nine classifiers (Table S1; Fig. S2a). We constructed a similarity network graph in which nodes represented classes and edge weights reflected the frequency at which samples were assigned to both classes connected by that edge (Fig. S2b). Edge weights between classes from the same classifier were zero. After network optimization (see *Methods*, Fig. S2c), we performed Markov clustering on the class network ^14^. Eleven classes from six classifiers resolved into two significantly interconnected clusters of classes (*p* = 2.6e-6, hypergeometric test, Fig. 1c) suggesting high overlap between existing classifiers. We named the clusters consensus cancer subtype-A and -B (CCS-A and -B).

Next, we developed a top scoring pair model^15^ for single-sample expression-based CCS assignment (eCCS; Fig 1d). We found relative expression at 16 pairs of genes could effectively distinguish samples according to CCS (Table S2; Fig. 1e). Testing the eCCS model in a held-out test set achieved 100% accuracy (95% C.I.: [84 - 100%]). Given the heterogeneity of the training data set, we also explored the impact of prior treatment and metastasis on model training; subsetting our data to only include treatment-naïve specimen or primary tumors, we only found disagreement in CCS assignment between 8 samples (6%; Fig S2d, e). While most samples had agreement in classification between all or most of the 16 gene pairs, these 8 samples had subtype assigned by a margin of only 1-2 gene pairs. Re-evaluation showed unusual clinical or genomic characteristics for 6 of the 8 cases (Table S3). Hereafter, we labelled samples assigned CCS by margins of less than three pairs as “Other” and removed them from analyses unless otherwise specified.

To extend analysis to datasets without gene expression data, we also trained a genetics-based model (gCCS) to predict CCS based on median relative arm-level copy number (*i.e.* log tumor / normal read counts; Table S4). This metric was chosen to allow for the use of the gCCS classifier on genomic data that do not cover the whole genome, such as exome or targeted sequencing. On a held-out test dataset (*n* = 30), a model predicting gCCS had an area under the receiver operator curve of 0.95 (Fig S2f). Validating the gCCS model on the data of Farshidfar *et al.* ^16^, the area under the curve was 0.99 (sensitivity = 1 where specificity = 0.79; Fig S2g).

### Consensus subtypes retain features of original classifiers

Three classifiers ^8–10^ contributing to CCS were specifically trained on intrahepatic cholangiocarcinomas. Exploring the alignment of anatomical location with CCS, we found eCCS-A samples appeared exclusively in intrahepatic cholangiocarcinomas or combined hepatocellular-cholangiocarcinoma, while eCCS-B spanned the biliary tract (Fig. 2a). The site of biopsy from metastases did not affect the association between eCCS and primary site of the tumor: all 23 liver metastases from extrahepatic primaries were assigned to eCCS-B. Analyzing 14 cases with both a primary and metastatic sample, all pairs exhibited concordant eCCS assignment despite different biopsy sites (CCS-A *n* = 9; CCS-B *n* = 4; Other *n =* 1). Within intrahepatic cholangiocarcinomas, the classifier from Song G. *et al*.^10^ was trained to distinguish between small and large duct pathologies. Using the cohort of Jolissaint *et al.* ^17^, the gCCS model recovered an enrichment of large duct intrahepatic cholangiocarcinoma within gCCS-B (*n* = 12, *p* = 0.01), while the small duct histopathology was significantly enriched in gCCS-A (*n* = 117, *p* = 0.01).

**Fig. 2.**
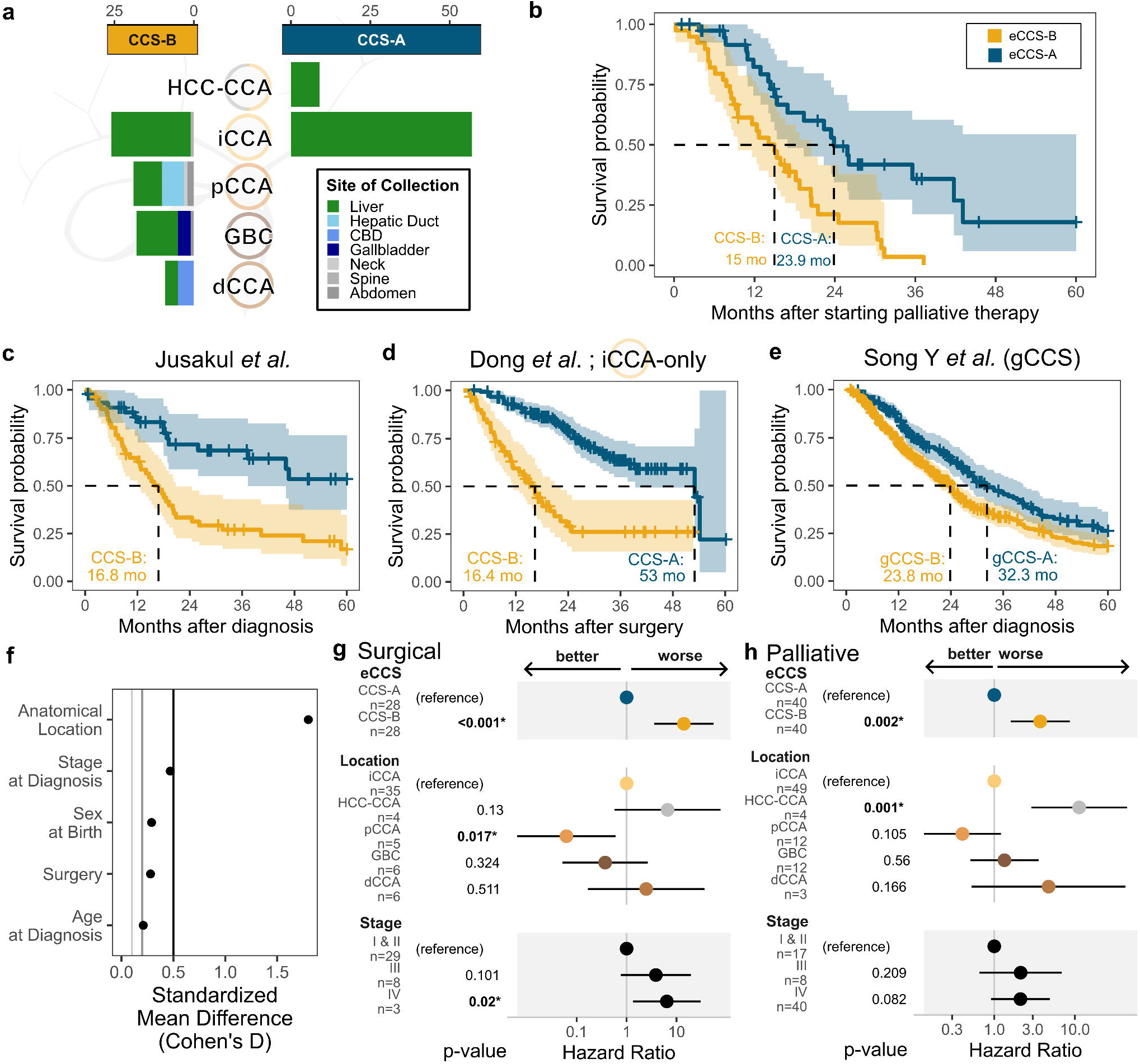
Survival differences between consensus cancer subtypes. **a**, counts of tumors classified by expression consensus cancer subtype (CCS) into anatomical location of primary tumor on the biliary tract, as well as site of specimen collection. **b**, overall survival after start of first-line palliative treatment by expression CCS in the present cohort. **c-e.** Kaplan-Meier analysis of survival **c,** from diagnosis in n = 97 BTC case from Jusakul *et al*. (2017) **d,** from surgery in n = 208 iCCA cases from Dong *et al.* (2021), and, **e,** using gCCS after diagnosis in n = 592 cases from Song Y *et al*. (2024) **f**, standardized mean difference in clinical covariates between the subtypes **g-h**, Cox proportional hazards models for eCCS, primary tumor location, and stage in **g**, surgical and **h**, palliative contexts. eCCS, expression consensus molecular subtype. CCA: cholangiocarcinoma; combined hepatocellular-cholangiocarcinoma (HCC-CCA), iCCA: intrahepatic CCA; pCCA: perihilar CCA, dCCA: distal CCA, GBC: gallbladder cancer. mo: months.

Two of the classifiers ^7,8^ were trained to maximize survival differences among patients. We found eCCS-A was associated with longer overall survival after surgery (*p* = 0.03) and after initiation of first-line palliative systemic therapy (*p* = 0.0004) than eCCS-B (Fig 2b). Assigning eCCS to tumor samples from external cohorts, patients with CCS-A tumors had significantly longer survival from diagnosis for 97 BTC patients from Jusakul *et al.* ^18^ (*p* = 0.0002; Fig 2c), which were assayed by array-based expression profiling. In the 208 intrahepatic cholangiocarcinoma patients from Dong *et al.* ^19^, CCS-A cases showed longer survival after surgery (*p* = 2.7e-11; Fig 2d). Stratifying patients from Song Y. *et al.*^20^ by gCCS using their genetic panel data (n = 592), we saw significantly longer overall survival in gCCS-A (*p* = 0.002; Fig 2e).

Given the convergence of CCS onto both an anatomical location and a survival difference, we tested whether anatomical location or CCS better predicted overall survival. We found no significant difference between CCS in commonly prognostic clinical variables including stage at diagnosis (*p* = 0.063), age (*p* = 0.4), sex (*p* = 0.2), or rate of resection (*p* = 0.4; Table S5; Fig 2f). In a multivariate Cox proportional hazard model including disease stage, eCCS was significantly associated with survival time after surgery (*p* = 3e-5) and after start of palliative treatment (*p* = 0.003; Fig. 2g, h), while most anatomical locations were not. Similarly, in the data from Song Y. *et al.*^20^, gCCS was significantly associated with survival time (*p* = 0.005), while anatomical locations were not.

### Subtypes reflect different tumor intrinsic transcriptional programs

eCCS groups 28% (*n =* 25) of intrahepatic cholangiocarcinomas with extrahepatic cholangiocarcinomas, suggesting higher-order biological structure not captured by anatomy alone. We tested whether grouping tumors by CCS explained more variation in gene expression than anatomical location. In a redundancy analysis, primary location and eCCS both explained significant variance in the first and third principal components of gene expression (*p* < 0.001), however eCCS explained 1.67-fold more variance in gene expression than anatomical location of primary tumor (Fig. 3a; S3a). Collapsing distal and perihilar cholangiocarcinomas into the broader category of extrahepatic cholangiocarcinoma, eCCS explained 1.69-fold more variance. We similarly saw more variance in methylation explained by eCCS in data from Jusakul *et al.*^18^ (Fig. 3b). These results show CCS captured more transcriptional variation than anatomical location.

**Fig. 3.**
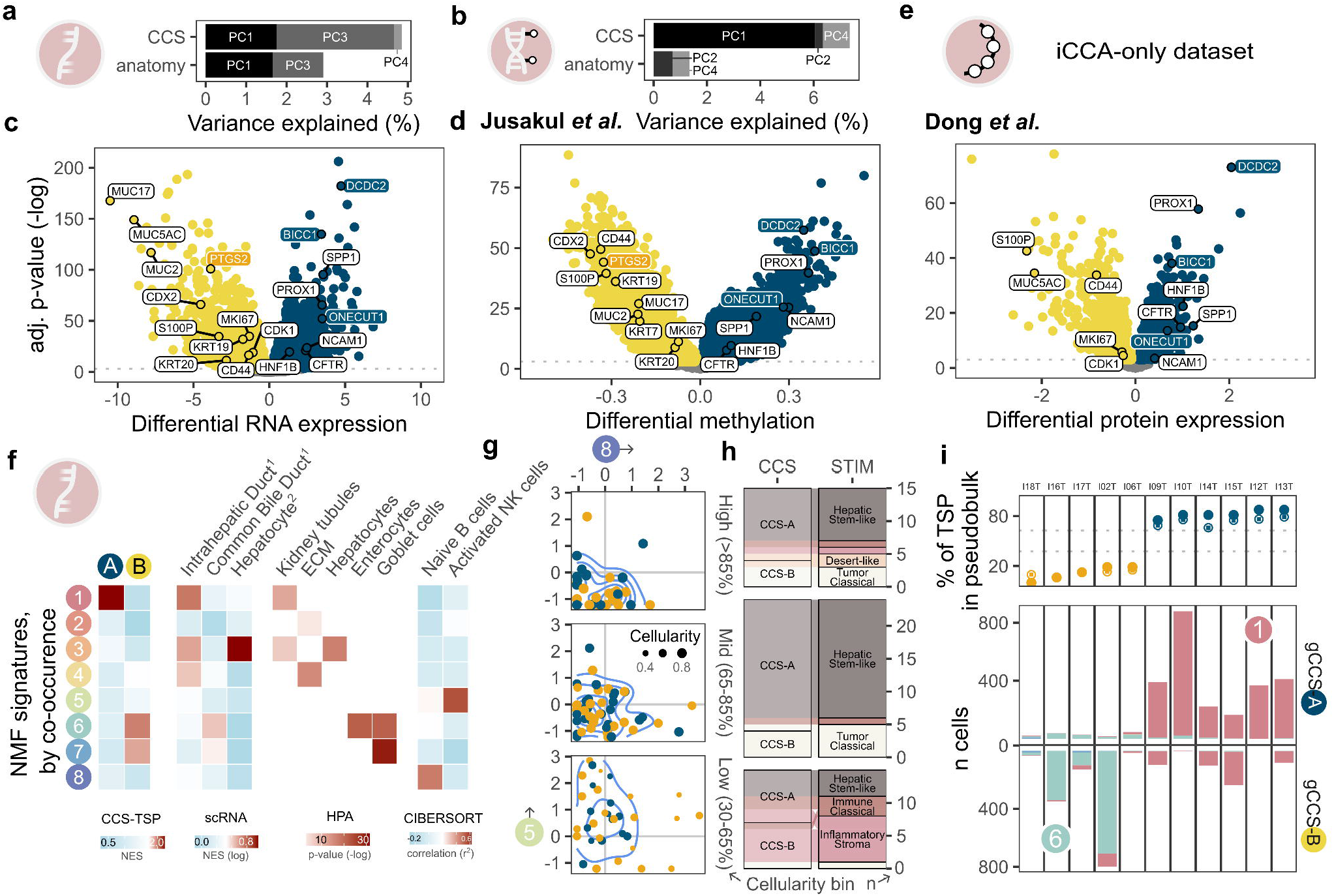
Differential expression and in silico microdissection of biliary tract cancer. **a-b,** comparing principal components of variance in gene expression across tumors as r^2^ of constrained variance in a redundancy analysis for location and consensus cancer subtype (CCS) in **a**, gene expression and **b,** methylation from Jusakul *et al*. (2017) **c-e**, differential **c,** gene expression, **d** methylation and **e,** protein expression between subtypes using the datasets from this study, Jusakul *et al.* (2017) and Dong et *al.* (2021) respectively. TSP genes in respective CCS colors. **f**, cell population marker enrichment for expression components from non-negative matrix factorization, including scRNA markers from ^1^Sampaziotis *et al.* (2021) and ^2^Andrews *et al.* (2024). **g-h,** Variance in tumor immune microenvironment across specimen tumor content using **g,** non-negative matrix factorization (NMF) gene expression signatures 5 and 8 and **h,** the Stroma-Tumor-Immune Microenvironment (STIM) classification of Martin-Serrano *et al.* (2023) across cellularity bins. **i**, eCCS per tumor by pseudobulk TSP, and tallies of cells by single cell genetic CCS (gCCS), with color by NMF signature. PC: principal component. TSP: top scoring pair.

Next, we enquired whether CCS reflects tumor intrinsic features or the tumor microenvironment. The top scoring pairs identifying eCCS-A include canonical cholangiocyte markers (*DCDC2*, *BICC1*, *ONECUT1*) suggestive of a signal from biliary epithelial cells. While genes from top scoring pairs for eCCS-B did not help resolve signal identity, differential gene expression showed eCCS-B tumors had higher expression of gastrointestinal epithelium markers (*CDX2*, *KRT19*, *KRT20*, *CD44;* Fig 3c.), which are also used to define intraductal papillary neoplasms of the bile ducts ^21^. Marker enrichment profiles were replicated in methylation data from Jusakul *et al.*^18^ and protein expression data from Dong *et al.*^19^ (Fig 3d, e). The enrichment of epithelial markers across data modalities supports that CCS reflect different epithelial-derived tumor cell populations.

Because bulk data collapses expression profiles from multiple cell types, we performed virtual microdissection using non-negative matrix factorization (NMF) to separate expression signatures. Varying the expected number of signatures, eCCS remained a top differentiating factor (Fig. S3b). When optimizing for components that correlated with tumor content, the data were best explained by eight signatures across 6,000 genes (see *Methods*; Fig. S3c).

Gene set enrichment analysis found two NMF-derived signatures (signatures 6 and 7) were significantly enriched for eCCS-B top scoring pair markers (false discovery rate [FDR] = 5e-6 and 7e-5, respectively; Fig. 3f). Both signatures were also enriched for scRNA markers of common bile duct epithelium ^22^ (FDR = 1e-9 and 2e-8) and genes corresponding to goblet cell proteins from the Human Protein Atlas ^23^ were significantly over-represented (FDR = 5e-10 and 4e-14). Goblet cells can appear in areas of the bile ducts and gallbladder associated with intestinal metaplasia ^24^. With the top scoring pair markers for eCCS-A, gene set enrichment analysis found one significantly enriched signature: signature 1 (FDR = 2e-9). Gene set enrichment analysis of scRNA markers for signature 1 found enrichment of intrahepatic duct markers (FDR = 7e-10), and, of Human Protein Atlas markers, we found overrepresentation of genes with protein expression in renal proximal tubules (FDR = 9e-6). Signature 3, with hepatocyte marker enrichment (FDR = 4e-8), was more common in low tumor content liver specimen as well as HCC-CCA cases (Fig S3d, e). Weights for signatures 1, 6 and 7 were highly correlated between primary and metastatic paired patient samples (Fig S3f). Associations between epithelial tissues and CCS-marker signatures were robust to cohort heterogeneity (Fig S3g).

NMF also uncovered two immune signatures. Weights for signature 8 were correlated with transcriptome-derived estimates of naïve B cells, resting CD4+ T cells, and resting NK cells (see *Methods*), suggestive of an immunologically quiescent environment. Signature 5 was associated with CD8+ T cell and activated NK cells suggestive of immune hot tumors. Specimens with low tumor content (<65%) showed larger NMF weights in immune signatures than specimens with high tumor content (>85%), where immune signal was mostly absent (Fig. 3g). Similarly, mapping intrahepatic CCA samples to the stroma-tumor-immune microenvironment (STIM) classification framework of Martin-Serrano *et al.*^25^, we saw a strong impact of tumor content on class assignment (Fig. 3h): in mid and high tumor content tumors, eCCS-A tumors map to the ‘hepatic stem-like’ class while eCCS-B tumors map to the ‘tumor classical’ or ‘desert-like’ classes, while in low tumor content specimens, eCCS-A tumors mapped to the ‘immune classical’ class and eCCS-B tumors were assigned to ‘inflammatory stroma’. Supporting our bioinformatic estimate of tumor content, we found a significant correlation between pathologist estimates of tumor content and bioinformatic estimate (r^2^ = 0.69; *p* = 0.001; Fig. S3h). Together, the NMF results suggest CCS, hepatocytes and immune factors load on distinct gene sets and are largely separable.

We next used single cell RNAseq from Song G. *et al.* ^10^ to confirm that CCS signal originated from individual tumor cells. We first inferred aneuploidy for each cell, a marker of malignancy ^26^, and used the inferred copy number profiles to predict gCCS for individual cells. Because gene dropout made eCCS assignment by top scoring pairs unreliable at single cell resolution, we projected the richer NMF-derived gene expression signatures onto single cells and reserved eCCS assignment for pseudobulked tumors. Tallying cells by their top NMF signature ^27^, we found most gCCS-B cells showed high levels of signature 6 and came from pseudobulked eCCS-B tumors, while gCCS-A cells upregulated signature 1 and appeared in pseudobulked eCCS-A tumors (Fig 3i). Quantifiably similar conclusions could be made with the scRNA BTC datasets from Zhang *et al.* ^28^ and Shi *et al.* ^29^ (Fig S3i, j). These results suggest CCS captures tumor cell-intrinsic transcriptional states.

### Subtypes harbor distinct genomic drivers

Studies of co-alteration patterns of mutations in pan-cancer driver genes have found recurrent genetic profiles in BTC ^4^ but previous studies have been underpowered to define rare BTC-specific drivers^30,31^. To test whether the CCS framework aligns with previously identified genetic profiles and to cement these mutations as statistically supported drivers, we turned to our whole genome sequencing for all 180 samples, calling BTC drivers *de novo*.

Eleven genes were significantly enriched for small substitution mutations (< 2 codons in length) across both a protein coding-centric and a coding-agnostic approach (see *Methods*), with two additional genes only passing the FDR in one approach (Fig. 4a, Table S6, S7A). The three genes with the strongest signal of over-representation across both tests were *TP53* (*n =* 62), *ARID1A* (*n* = 33) and *BAP1* (*n =* 31). Among gene promoters, we identified recurrent small substitutions in *TERT* (*n* = 8, FDR *q* = 0.02).

**Fig. 4.**
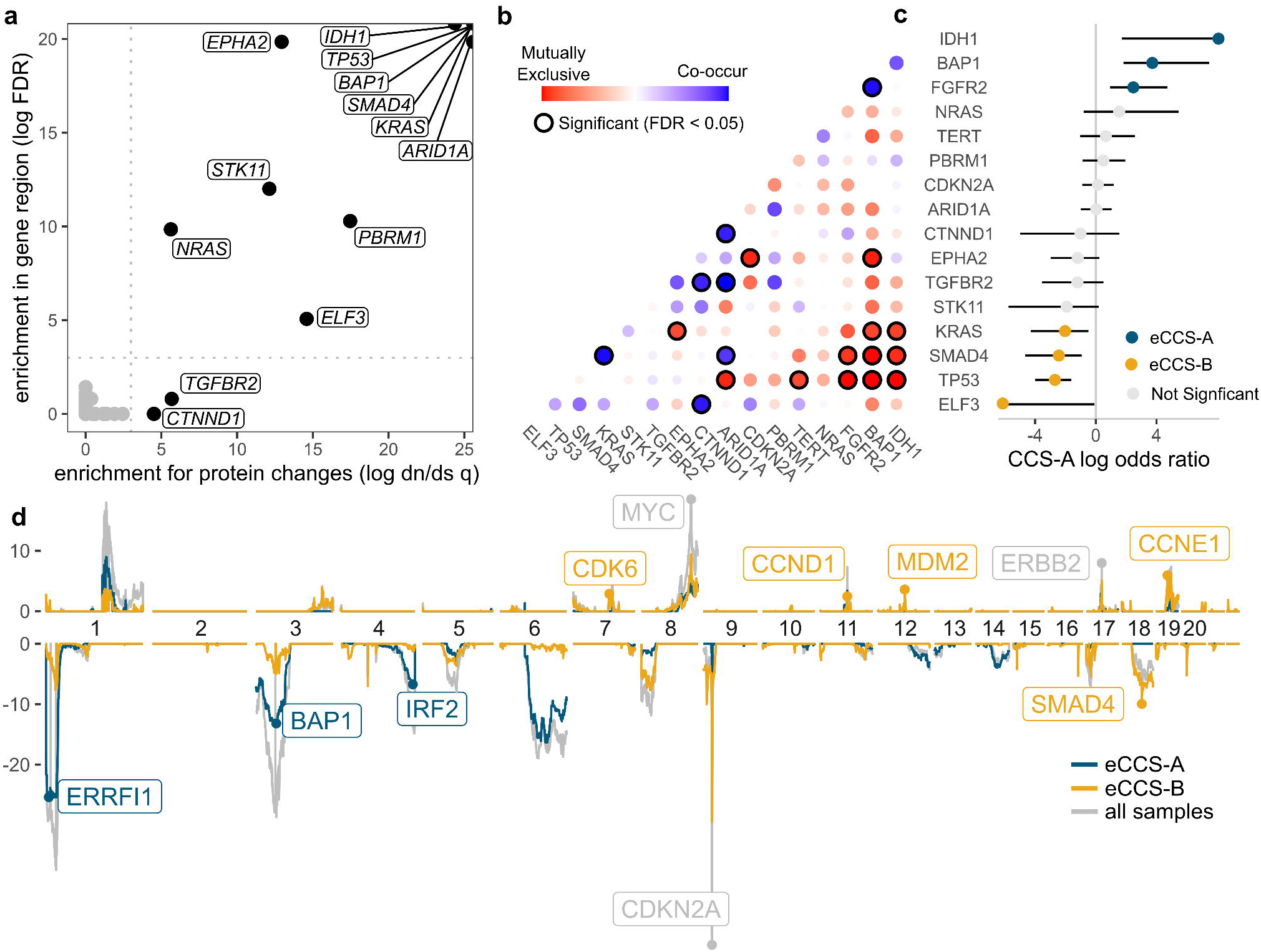
Genetic drivers of biliary tract cancer. **a**, statistical recurrence analysis for small mutation drivers for a gene-centric approach (the rate of non-synonymous to synonymous mutations, dn/ds) and gene-agnostic approach (regional enrichment). **b**, co-occurrence matrix for drivers, with enrichment for co-occurrence in blue and mutual exclusion in red, and significant associations circled in black. **c**, Log of the odds ratio for driver mutations by consensus cancer subtype (CCS). **d**, significance of copy number alteration recurrence using GISTIC, across chromosomes and color by CCS and using all samples (globally). FDR: False discovery rate.

Next, we explored genes impacted by recurrent structural variants (Table S7B). To enrich for functional impact, we tested structural variant breakpoints overlapping expression quantitative trait loci ^32^. Top structural variant expression quantitative trait loci were at *FGFR2* (*n =* 29), *TERT* (*n =* 3), *CDK5* (*n =* 8) and *MDM2* (*n =* 5). Amplicon reconstructions from copy number profiles found the regions encompassing *MDM2* and *CDK6* were commonly extrachromosomal DNA (*n* = 4 of 5 and *n* = 3 of 8, respectively). Because most high effect size structural variants were rare (<5% of cases), we formally assessed recurrence with a Q-Q enrichment analysis ^33^. *FGFR2* was the only gene with somatic structural variants showing significant upregulation and statistical recurrence (FDR = 9e-8).

We explored how mutations in drivers were co-distributed among tumors (Fig. S4a) and how this related to CCS. Testing how often samples were mutated versus wildtype, we recovered a nexus of mutual exclusivity defined by *TP53*, *SMAD4*, and *KRAS* mutations in eCCS-B cases, in contrast to mutations in *IDH1*, *FGFR2*, and *BAP1* in eCCS-A (Fig. 4b, c). Small substitution driver analysis restricted within each eCCS confirmed these findings in addition to recovering *CDKN2A* (*n* = 7, *q* = 1e-4) and *ARID2* (*n* = 10, *q* = 0.01) as drivers exclusive to CCS-B (Fig. S4b). Subsetting across axes of cohort heterogeneity suggested possible impacts of prior treatment and metastasis on rates of drivers within CCS but the CCS-defining drivers remained consistent (Fig. S4b).

Next, we examined copy number variants across BTC (Table S7C). Evaluating recurrence of copy number alterations ^34^, we found well-characterized oncogenes such as *ERBB2* and *MYC* as well as recurrent homozygous deletions of tumor suppressor gene *CDKN2A* (Fig. 4d). Analysis within CCS revealed divergent copy number profiles: recurrent narrow amplifications were over-represented in eCCS-B, including *CCND1* and *CCNE1*. eCCS-B tumors were also more likely to exhibit extrachromosomal DNA (Fisher’s exact test; *p* = 0.0006). In contrast, eCCS-A commonly showed chromosome-arm scale heterozygous deletions (single arm losses). Tumor suppressor genes in other cancer types with changes in gene expression in eCCS-A included *ERRFI1* and *IRF2*, but we also note recurrent deletion of 6q and amplification of 1p21 without conclusively identifiable drivers.

Under Knudson’s two-hit hypothesis ^35^, copy-loss events leading to loss-of-heterozygosity should converge on tumor suppressor genes. Six driver genes showed significantly more loss-of-heterozygosity events in mutated compared to wildtype cases, with loss-of-heterozygosity occurring at the highest rates with mutations in *TP53* (53 loss-of-heterozygosity events among 62 mutations, FDR = 1e-14; Fig. S4c). Loss-of-heterozygosity did not occur with mutated *IDH1* and *NRAS*, supporting their roles as oncogenes rather than tumor suppressor genes in BTC.

We posited that the segregation patterns of drivers could play a role in differences in survival outcomes between CCS. Because *TP53* is a prognostic factor in BTC ^36^, we tested whether eCCS was still prognostic in *TP53* wildtype cases. Subsetting our palliative cohort to *TP53* wildtype cases, we saw a trend towards longer survival in eCCS-A, but the difference was not significant (*p* = 0.07), potentially because only 20 eCCS-B cases remained. In the 171 *TP53* wildtype intrahepatic cholangiocarcinomas from Dong *et al.*^19^, eCCS was significantly associated with overall survival after surgery (*p* = 5e-9). Furthermore, in these data, a multivariable Cox proportional hazards model including eCCS, *TP53* status and tumor stage found a significant association with survival and eCCS (*p* = 4e-7) but not *TP53* (*p* = 0.66; Fig. S4d). Another possible factor confounding survival with CCS was the strong segregation of clinically targetable alterations in CCS-A, as both *FGFR2*-fusion and *IDH1*-mutated tumors have FDA-approved therapies. In our cohort, one case was treated with pemigatinib (an FGFR2-inhibitor), while four were treated with an IDH1-inhibitor on trial in the palliative setting. When these cases were removed from the analysis, we still observed a significant difference in survival by eCCS (*p* = 0.004).

### Divergent evolutionary dynamics between subtypes

Turning to the broader mutational context, we observed a 1.6-fold increase in tumor mutational burden (TMB) in eCCS-B over A (*p* = 3e-6, Fig 5a). This difference remained significant after excluding highly mutated (TMB > 10) samples (fold change = 1.4, *p* = 5e-5) and therefore is not solely explained by the numerical increase in rates of homologous recombination deficiency (*n* = 6 and 2) and microsatellite instability (*n* = 4 and 0) in CCS-B over CCS-A. eCCS was a better predictor of TMB than primary location: in a two-factor ANOVA, eCCS was significant (*p* = 0.002) while primary tumor location was not (*p* = 0.64).

**Fig. 5.**
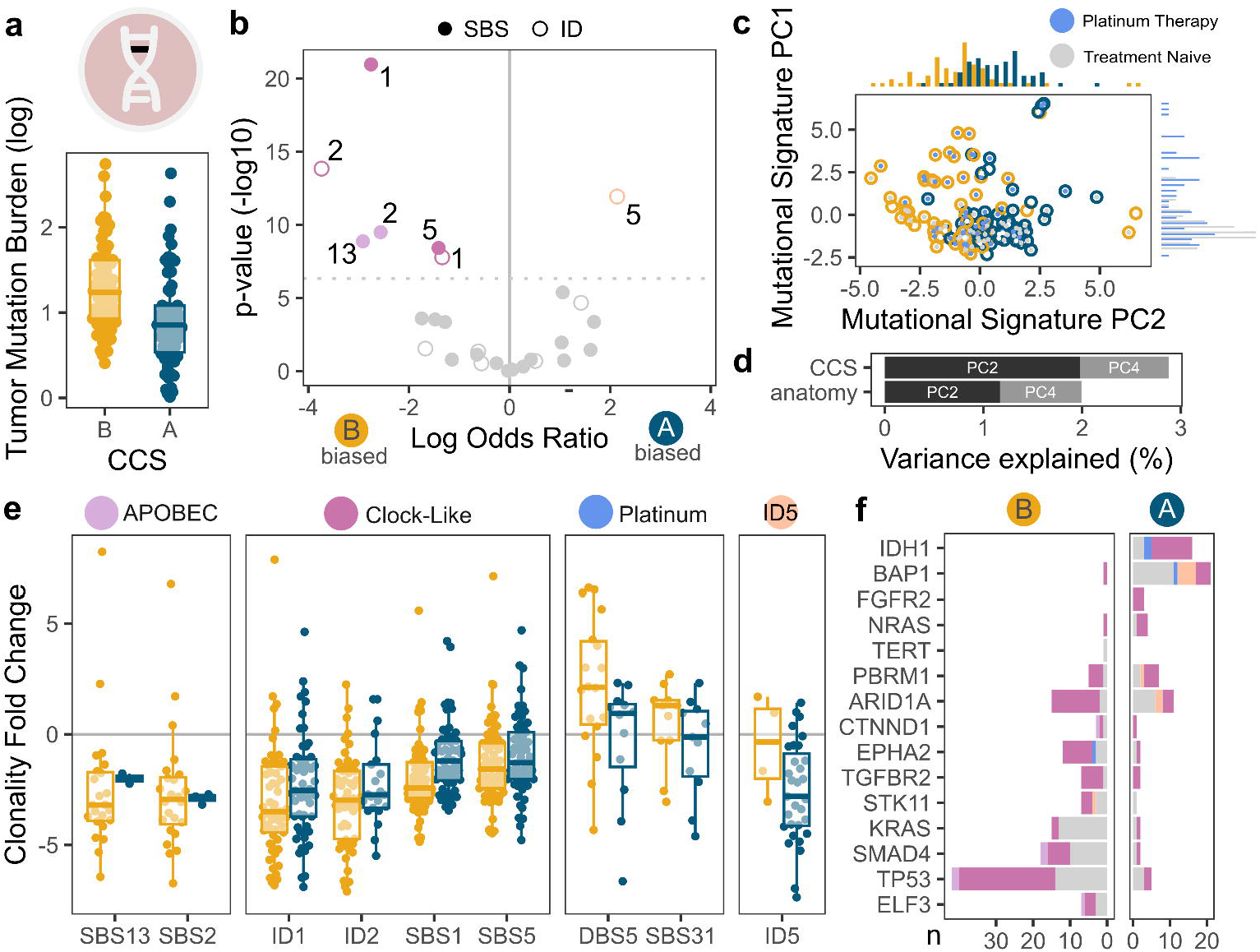
Mutation profiles of biliary tract cancers. **a**, tumor mutational burden between consensus cancer subtypes (CCS). **b**, enrichment in mutational signatures for single basepair substitutions (SBS, ●) and in/dels (ID, ○) by subtype. **c** principal component analysis of mutational signatures, color by platinum therapy naïve (grey) and exposed (blue) samples and CCS. **d**, variance explained by location and CCS in redundancy analysis of principal components of gene signature variance **e**, relative evolutionary timing of clonal mutations by mutational signature and CCS. **f,** mutations in genetic drivers by mutational signature and CCS. PC: principal component.

We next compared whether the increased burden could be explained by specific mutational signatures. *De novo* mutational signature extraction identified 22 single basepair substitution (SBS), seven double basepair substitution and eight in/del (ID) signatures (Fig. S5). Comparing signature contributions between CCS using a linear model, eCCS-B tumors showed significant enrichment of SBS1/SBS5 (clock-like; *p* = 8e-10), ID1/ID2 (clock-like; *p* = 9e-7), and SBS2/SBS13 (APOBEC; *p* = 7e-5 and 1e-4), whereas eCCS-A tumors showed enrichment of ID5 (unknown etiology; *p* = 6e-6; Fig 5b). Further, CCS-associated signatures were found to be a principal component of variance among samples. The second principal component of variance in mutation counts assigned to each signature, explaining 8.4% of the variance, distinguished eCCS-A from eCCS-B tumors (*p* = 3e-11; Fig. 5c). The first principal component (PC1), explaining 9.5% of the variance, loaded on SBS31 and DBS5, both associated with platinum therapy. As expected, patients who received platinum-based chemotherapy before specimen collection had significantly higher PC1 values (*p* = 9e-6). Once more, CCS better explained variance than primary location: redundancy analysis found eCCS explained 1.4-fold more variance than primary location. These results suggest not only significant differences in mutational profiles between CCS, but that these differences were among the top axes of diversity among samples.

Finally, we hypothesized that differences in mutational burdens might reflect tumor evolution. Among clonal mutations, signatures showing CCS-bias occurred early in that subtype’s evolution: relatively more APOBEC and clock-like signatures were earlier than later in eCCS-B (*p* = 6e-5 and *p* = 9e-12, respectively) with a similar finding for ID5 in eCCS-A (*p* = 6e-5; Fig. 5e). Further, these signatures contributed considerably as mutations in driver genes in their respective subtypes (Fig 5f). These results suggest these signatures may be important in early stages of subtype evolution.

## Discussion

Molecular subtyping has stratified many cancers into biologically and clinically meaningful groups. However, in BTC, classification has been anchored to anatomy or mutations in specific genes. In this study, we integrated published classifiers to define two subtypes, CCS-A and CCS-B. By integrating whole-genome and transcriptome sequencing in a large cohort, we demonstrate that CCS capture more variation across gene expression, driver mutations, mutational signatures, and clinical outcomes than anatomical classification. We demonstrated that the CCS subtypes reflect different tumor-intrinsic epithelial expression programs. The association between CCS-A and small duct single-cell markers and CCS-B with extrahepatic markers, combined with the distinct embryological origin differences between these tissues ^37^, suggest that the CCS may stratify BTC based on distinct cells of origin. We anticipate that larger BTC cohorts will reveal further stratification within the subtypes and expect stratification to exist within the tumor microenvironment which may be better captured by other technologies ^38^.

Whole genome sequencing supports divergent evolutionary trajectories between CCS. We identified drivers in CCS-B newly associated with BTC, including *EPHA2.* We also revealed putative tumor suppressors in CCS-A, including *ERRFI1*, which has potential therapeutic implications ^39^ and *IRF2* which emerged as a candidate gene for BTC predisposition in a genome-wide association study ^40^. We also found distinct copy number profiles between subtypes, with CCS-B enriched for narrow high-copy number changes. Extrachromosomal DNA provides a mechanism for oncogene activation in CCS-B, exemplified by amplification and overexpression of *MDM2*. CCS-B tumors have higher mutational burdens, mostly driven by clock-like or APOBEC-associated signatures, starting early in tumor evolution. The association between the enrichment of molecular clock signatures and BTC precursor lesions and metaplasia in CCS-B could provide fruitful insights in the early stages of evolution of this subtype. CCS-A tumor cells exhibited few well-characterized drivers and a lower mutational burden, suggesting its evolution may be less mutationally-driven than CCS-B. Multiple specimen from the same patient would resolve how these patterns translate to intratumoral heterogeneity.

The CCS framework provides an organizing system for BTC that balances the trade-off between grouping similar tumors to maintain statistical power in clinical trials for such rare cancers and distinguishing features of meaningful subtypes. CCS harmonizes across data modalities and shows inter-platform reproducibility but does not disqualify the worth of each classifier in its trained context. While our work suggests CCS is prognostic, we did not test whether CCS is a predictive biomarker. Because most patients received standard therapies such as adjuvant capecitabine and first-line chemoimmunotherapy, we were unable to evaluate potential predictive effects. However, we anticipate that accounting for CCS in clinical trials may be important for discerning outcome differences.

In conclusion, our data provide the first comprehensive integration of multi-omic and clinical evidence to define consensus cancer subtypes for BTC. Bridging previously disparate observations and infusing the field with the largest new genomics dataset in a decade, we lay the groundwork for a new era of precision oncology in this historically heterogeneous disease.

## Acknowledgements

This research was supported by the Richard LeGresley Biliary Cancer Research Fund through the Princess Margaret Cancer Foundation. Sequencing was supported by The Terry Fox Research Institute (TFRI) Marathon of Hope Cancer Centres Network (MOHCCN). We thank all patients and caregivers who contributed to this research. The authors used ChatGPT for recommendations for improving code in simple functions, and for suggesting improvements to manuscript flow and consistency.

