## Supplementary Materials for "Integrated whole-genome and transcriptome sequencing reveals divergent evolutionary processes across biliary tract cancer subtypes"

### Supplementary Methods

*Case annotation*

All clinical data were reviewed according to the LeGresley data dictionary and logged in Redcap. Surgery was marked as aborted if surgery was attempted but aborted intraoperatively due to metastatic disease (planned procedure not completed/aborted).

*Sample processing and sequencing*

A unique tumor sample, from the primary lesion or a metastatic site, with paired normal tissue was included per patient. Samples then underwent whole genome sequencing, prioritizing fresh frozen laser capture microdissected samples), achieving a median coverage of 85-fold for the tumor, and 38-fold for the normal. Tumors also underwent whole transcriptome sequencing (WTS), when enough tissue remained, with a median of 83 million reads per sample.

Whole-genome and transcriptome libraries were prepared and Illumina sequencing was performed by the Ontario Institute of Cancer Research. Sample processing is previously described in Beaudry *et al* [12]. Briefly, whole genome sequencing fastq files from each lane were adapter trimmed using *cutadapt* 1.8 [62] and aligned to hg38.p14 using *bwa mem* 0.7.17 [63]. Optical replicates were marked by *Picard* 2.21.4 and lanes collapsed into a single .bam file. Median coverage was estimated with *Picard CollectWgsMetrics,* to a minimum of 80X for tumor samples and 30X for normal samples; additional sequencing was performed when this threshold was not met. For estimating tumor content, B-allele frequencies were derived from germline single nucleotide variants called using GATK's v. 4.1.2 *haplotypecaller* [64], with calls recalibrated and realigned using *Picard*. Read depth was segmented and integrated with B-allele frequencies using *HMMCopy* 0.1.1 and copy number alterations, tumor content and ploidy were called using *Celluloid* v0.11.7 [40]. All *celluloid* solutions were manually reviewed and samples estimated to have less than 30% cellularity were excluded.

Small substitutions including single basepair substitutions (SBS), double basepair substitutions (DBS) and insertion/deletions (in/dels) were called using GATK *mutect2* and *strelka2* 2.9.10 [65] and combined on consensus of both callers. Structural variants were called using *SVaBa* 1.2.0 [66], *delly2* 1.0.3 [67] and *manta* 1.6.0 [68], and combined on consensus of at least two of the three callers. Extrachromosomal DNA was inferred using *AmpliconArchitect* 1.3.3 [69], downsampling to 8 reads (--downsample 8) and copy number gain threshold set to 8 (--cngain 8); runs where a solution was not reached in 20 days of runtime were excluded from ecDNA analyses ($n$ = 6).

Tumor mutational burden was calculated as the sum of all small substitutions divided by genome size (in megabases). Microsatellite instability was called using *msisensor-pro* 1.2.0 [70] thresholding MSI-H as greater than 15% of sites showing instability. Candidate homologous recombination deficiency cases were identified using *HRDetect* from *signature.tools.lib* 2.2.0 [71,72] with a cutoff of 90% probability of HRD and reviewed manually.

Clonality was called using *pyclone-VI v* 0.1.6 [73]. Mutations were timed using the mutationTime function with n.boot = 10 and rho = 0 in *MutationTimer* v 1.0.2 [74]. Signature timing was tested by comparing between the distributions in relative exposures of clonal early and clonal late in a Wilcoxon rank-sum test.

RNA reads were mapped to hg38.p14 using *star* 2.7.4a [75]. Fusions were identified using *star-fusion* 1.9.0 and annotated with *mavis* v2.2.9 [76]. Reads were tallied per transcript using *htseq* v2.0.4 [77]. Whole transcriptome samples where the median reads per gene was less than one were excluded. Principal component analysis was performed with *FactoMineR* v. 2.12.

All analyses in R were performed on version 4.5.1 and in python on version 3.10.6.

*Processing public datasets*

For testing in external datasets, RNA data from Dong *et al* [19] was extracted from Table S3 and proteins from Table S5 of that manuscript. RNA beadchip data from Jusakul et al [18] was downloaded from cbioportal.org and normalized with *lumi* v 2.60.0 [78], and methylation data was downloaded from GEO Series GSE89803 and normalized with *minfi* v 1.54.1 [79]. Survival differences between the CCS were evaluated in a cox proportional hazard model including overall stage at diagnosis and anatomical location of the primary when appropriate (samples from Dong *et al.* were all intrahepatic) using *coxph* from *survival* v 3.8.3.

For public single cell RNAseq data analysis, data from Zhang et al [28] were downloaded from GEO Series GSE138709, from Shi et al [29] from GSE201425 and data from Song G et al [10] were obtained by personal communication with study authors. Data were normalized using *SCTransform* in *Seurat* v 5.3.0 [80]. We used markers from CellMarker [81] to define cell populations, subsetting to cell populations with epithelial markers or where marker-based cell identity was inappropriate for the context (e.g. acinar cells in the liver) as candidate malignant cell populations. NMF signatures were projected onto single cells by selecting the genes with the top 90th percentile of weights in their respective signatures, and testing for area under the curve-based enrichment with *AUCell_calcAUC* from *AUCell* v 1.30.1 [27]. Copy number alteration profiles were inferred from *infercnv* v 1.24.0 [26] per sample, using immune cell populations as reference and including a denoising step (denoise=TRUE). eCCS call confidence was estimated by resampling 100 subpopulations of 20 cells per tumor sample.

### Supplementary Figures

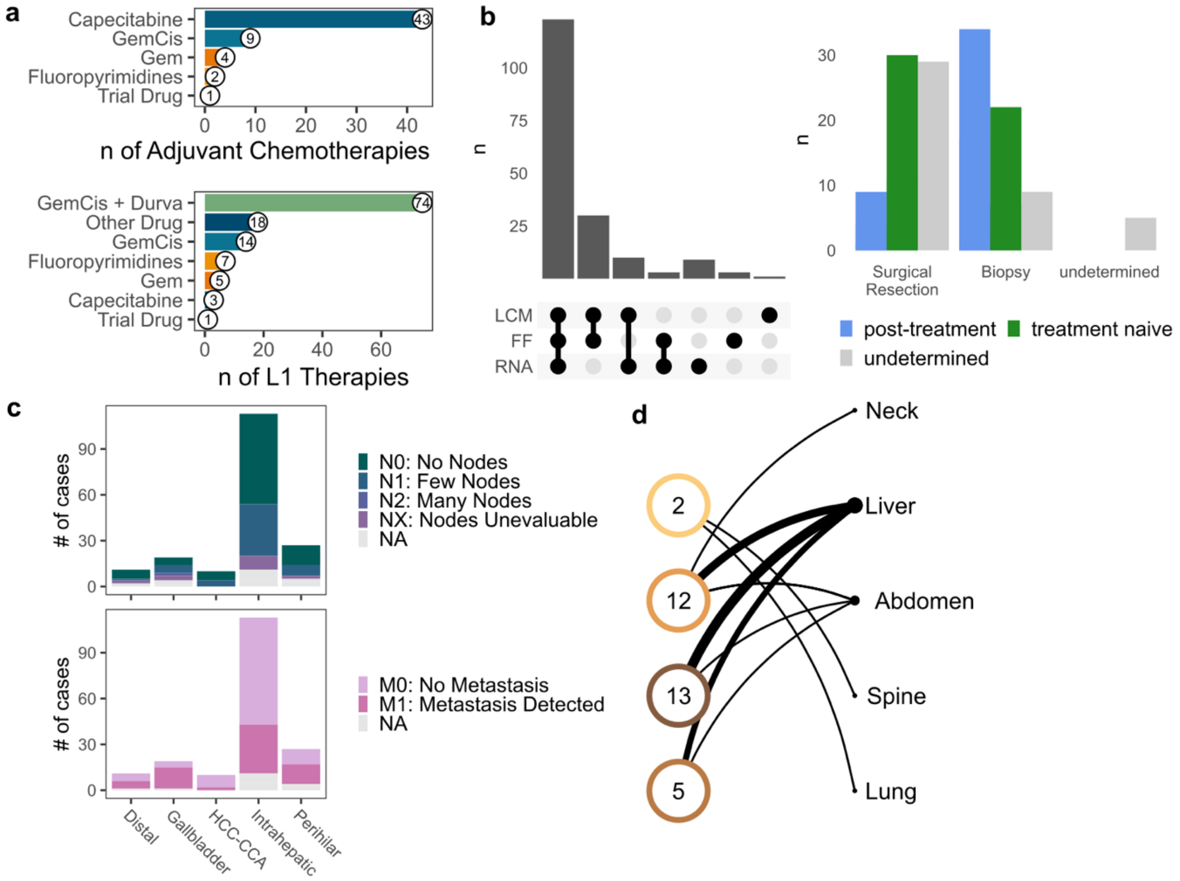


**Figure S1**

**Fig. S1 | Cohort specimens**

**a,** counts of patients by adjuvant chemotherapies and first line therapy **b,** counts of specimens by collection features including paired RNA (WTS), laser capture microdissection (LCM), and fresh frozen (FF) as opposed to formalin-fixed paraffin embedded (FFPE); most samples are LCM fresh frozen samples with paired RNAseq (WTS) as well as counts of specimens from biopsy compared to surgical resection, by timepoints of treatment naive (green) and post-treatment (blue). **c,** node (N) and metastasis (M) status across primary tumor locations **d**, Counts of biopsy sites of metastatic samples.

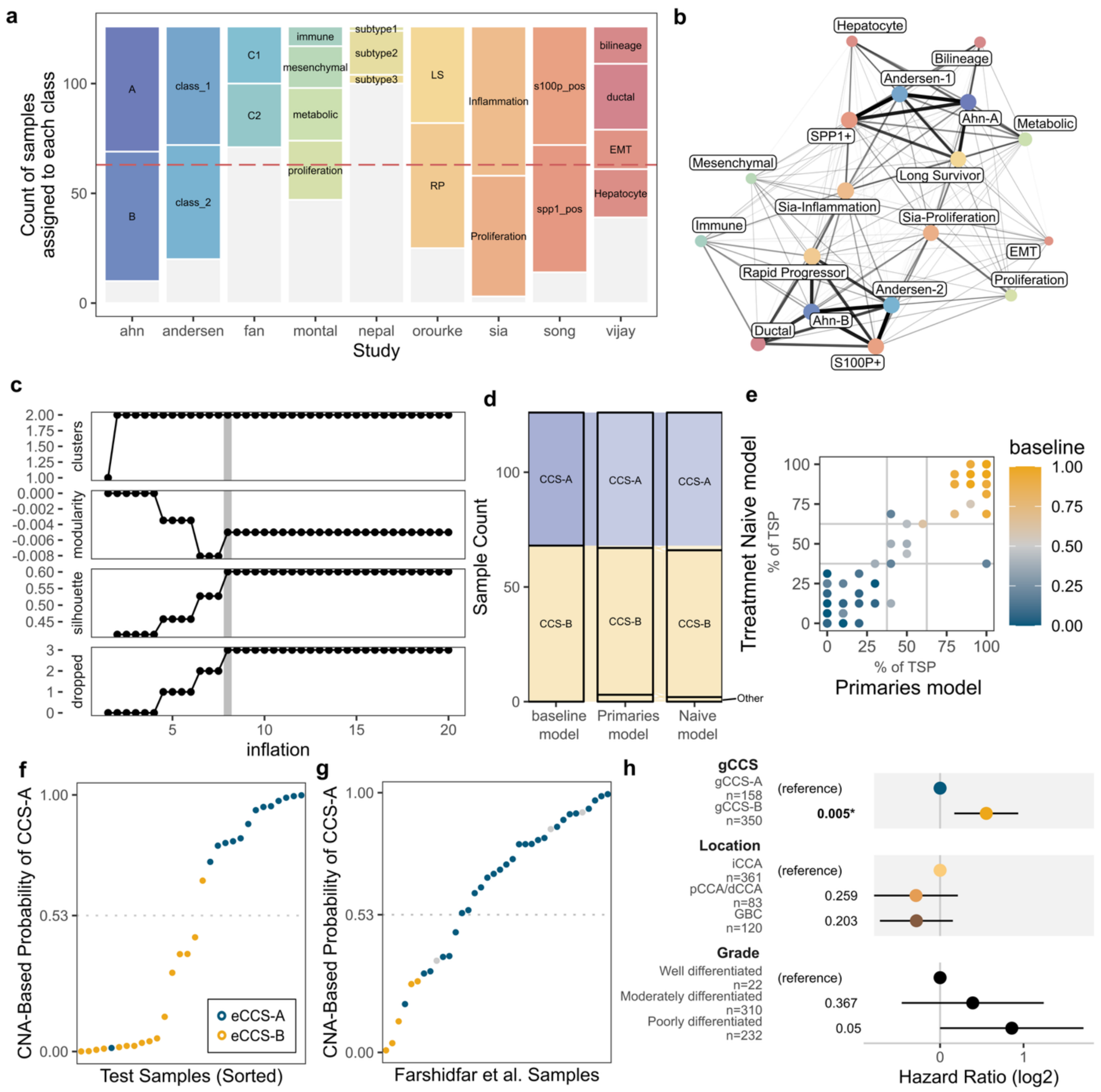


**Figure S2**

**Fig. S2 | eCCS training cohort and external validation**

**a,** assignments of 126 fresh frozen RNA samples to respective classes from nine previously published classifiers. Classifiers to have greater than 50% of cases assigned to any class (dashed line) to be included in network. **b,** expression classifier network before optimization. **c,** four parameters used for network optimization across inflation factors: number of clusters, modularity, silhouette score, and number of classes dropping out of the network. The gray band shows the chosen optimal inflation factor. **d,** CCS sample assignments in classifiers trained on data subset to only primary tumors (primaries model) and treatment-naïve tumors (Naïve model). **e,** Agreement among top score pairs in CCS class for the primary and treatment-naïve models, with the baseline model as point color TSP agreement. **f-g**, Accuracy of copy number alteration generalized linear model at predicting expression CCS (eCCS) for **f,** a test cohort (n = 30) and **g,** data from Farshidfar *et al.* (2017) **h,** Cox proportional hazards for genomic CCS (gCCS), location and grade on overall survival from diagnosis for unresectable disease for 592 cases from Song Y. *et al* (2024).

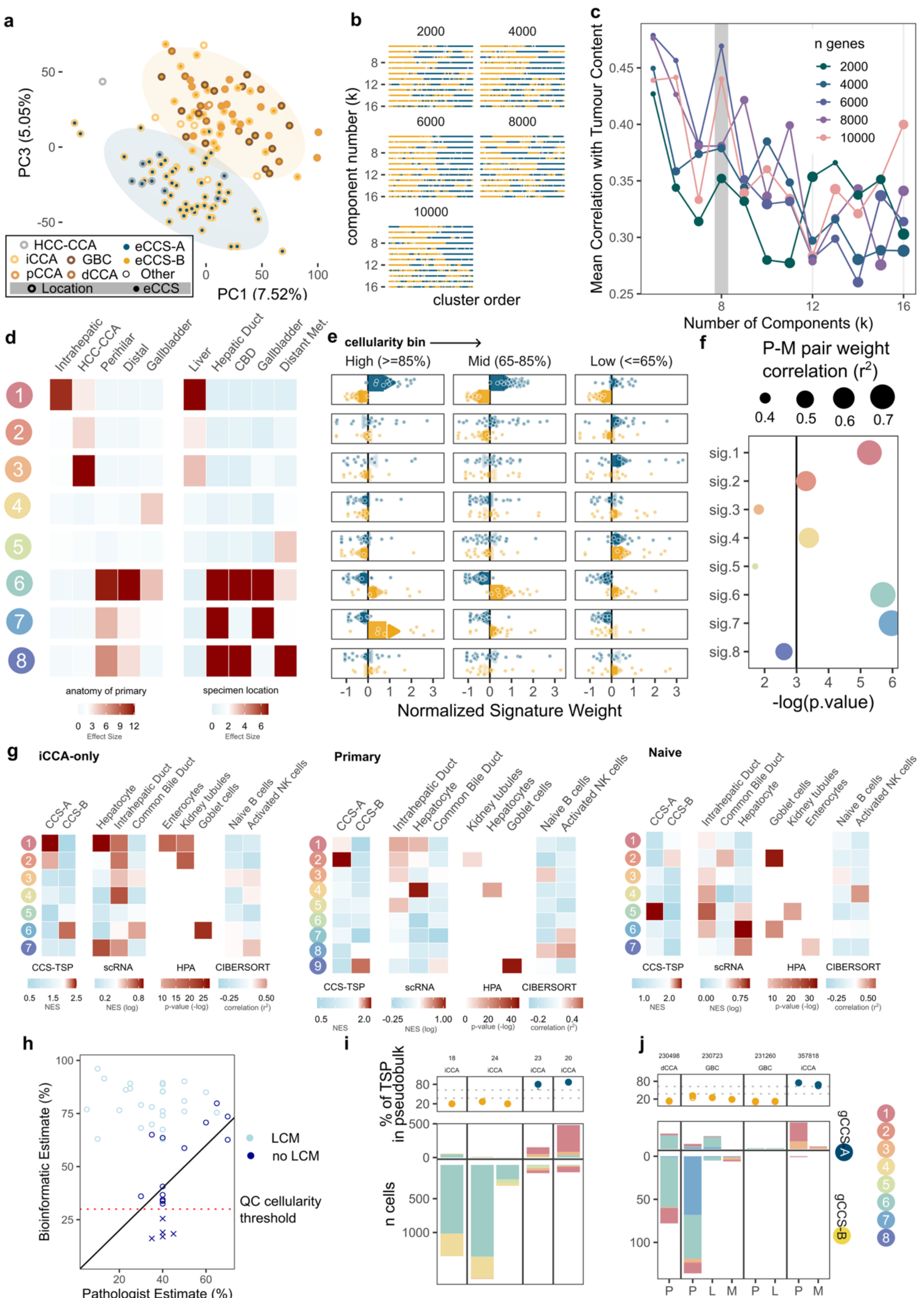
a
PC3 (5.05%)
PC1 (7.52%)
HCC-CCA
iCCA
pCCA
GBC
dCCA
eCCS-A
eCCS-B
Other
Location
eCCS
b
component number (k)
cluster order
2000
4000
6000
8000
10000
c
Mean Correlation with Tumour Content
Number of Components (k)
n genes
d
anatomy of primary
specimen location
Effect Size
e
cellularity bin
High (>=85%)
Mid (65-85%)
Low (<=65%)
Normalized Signature Weight
f
P-M pair weight correlation (r²)
-log(p.value)
g
iCCA-only
Primary
Naive
CCS-TSP
scRNA
HPA
CIBERSORT
h
Bioinformatic Estimate (%)
Pathologist Estimate (%)
LCM
no LCM
QC cellularity threshold
i
% of TSP in pseudobulk
n cells
Zhang et al. samples
j
Shi et al. samples
gCCS-A
gCCS-B

**Figure S3**

**Fig. S3 | Expression deconvolution and single cell RNAseq**

**a.** principal components of variance in gene expression across tumors (with inner color as primary location and outer color as eCCS). **b,** varying number of components and genes included in the non-negative matrix factorization with sample color by eCCS and clustered by hierarchical clustering**. c,** parameters for manual optimization of gene count (color) and component count (k) for non-negative matrix factorization of transcript per million (TPM) expression data, using mean correlation of all components filtered for a correlation with WGS-estimate of sample tumor content greater than 0.2. Vertical gray bar shows chosen optimal k. **d,** effect size of enrichment of presence of gene expression signatures derived from NMF by anatomic location of primary and location of specimen collected. **e**, normalized signature weights in CCS-A and -B specimen across cellularity bins. **f**, correlation in signature weights between paired primary and metastatic samples from the same case. **g**, de novo deconvolution of expression signatures using data subset to iCCA-only, primary tumors only, or treatment naïve tumors only. **h**, correlation between pathologist estimated tumor content and bioinformatics tumor content estimate for laser-capture microdissected (LCM) and non-LCM specimen. **i-j,** bootstrapped pseudobulk eCCS assignment with proportion of genes in TSP agreeing with call, and tallies of aneuploid cells by top NMF-expression signature assignment for scRNA of tumor samples from **i,** Zhang *et al.* and **j**, Shi *et al.*

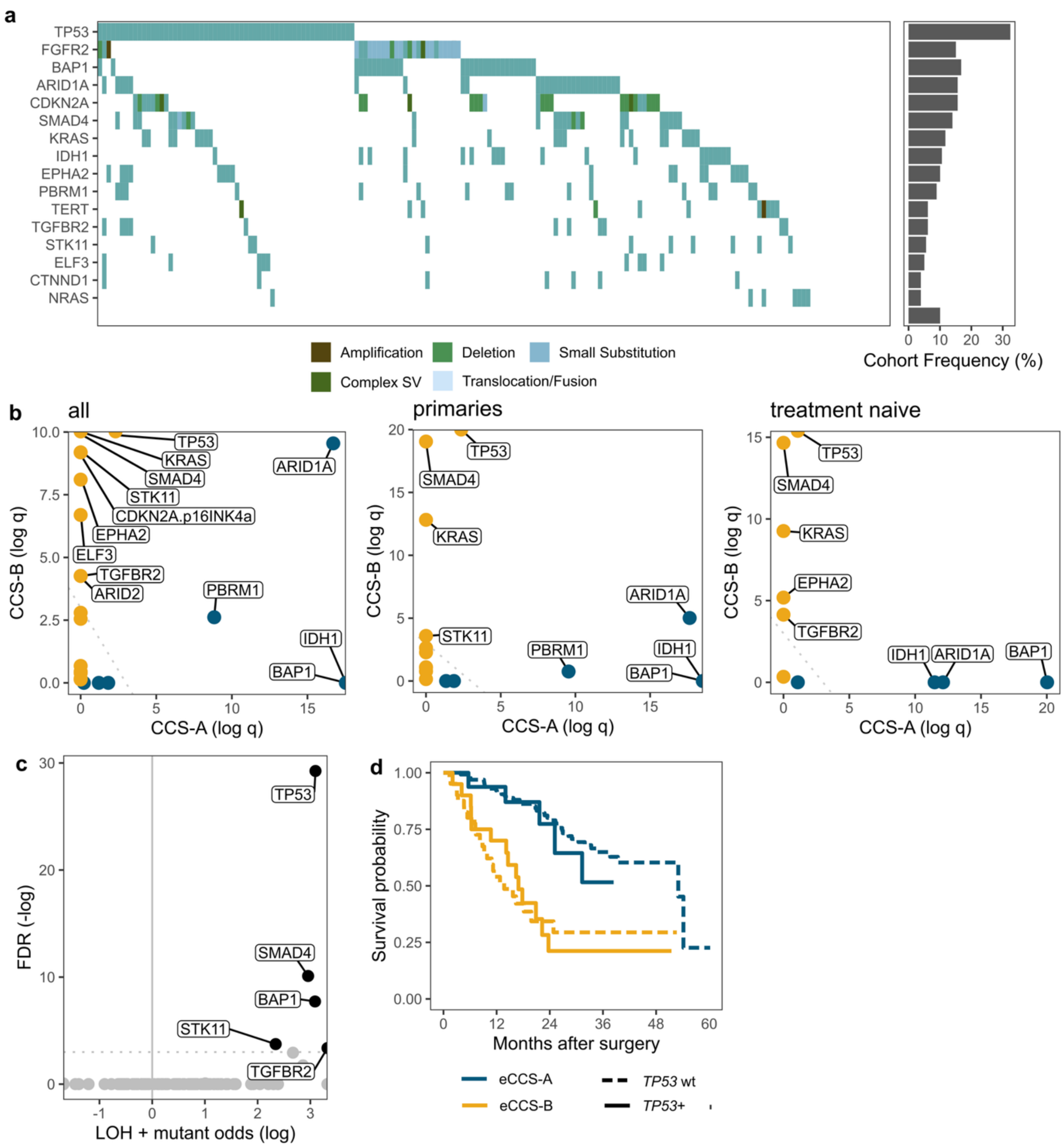

a
TP53
FGFR2
BAP1
ARID1A
CDKN2A
SMAD4
KRAS
IDH1
EPHA2
PBRM1
TERT
TGFBR2
STK11
ELF3
CTNND1
NRAS
Cohort Frequency (%)
Amplification
Deletion
Small Substitution
Complex SV
Translocation/Fusion
b
all
primaries
treatment naive
CCS-B (log q)
CCS-A (log q)
TP53
KRAS
SMAD4
STK11
CDKN2A.p16INK4a
EPHA2
ELF3
TGFBR2
ARID2
ARID1A
PBRM1
IDH1
BAP1
c
FDR (-log)
LOH + mutant odds (log)
d
Survival probability
Months after surgery
eCCS-A
eCCS-B
TP53 wt
TP53+


**Figure S4**

**Fig. S4 | Extended data on drivers by CCS**

**a,** oncoplot of small substitutions and structural variants across genes with significant mutational over-presentation, by mutation type; cohort frequency in right panel. **b,** genes showing significantly elevated dN/dS ratios in analysis within eCCS, as well as subset for only samples from primary tumors and for treatment-naïve tumors. **c,** over-representation of loss-of-heterozygosity in mutated cases for genes with significant mutational over-representation before multiple testing correction in previous analysis. **d,** Kaplan-Meier analysis of survival by eCCS in Dong. *et al.* by TP53 status.

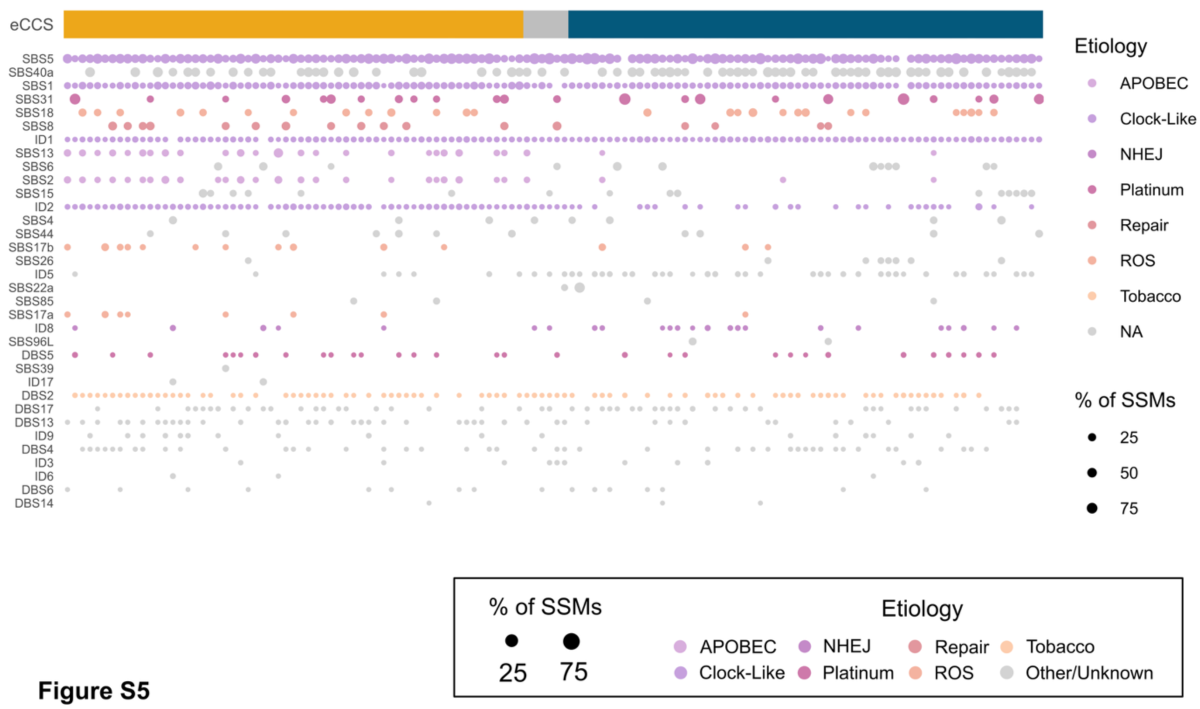


**Fig. S5 | Mutational signatures**

Full mutational signature assignments across samples and signatures, size as proportion of mutations in sample, clustered by mutational profile, top panel showing eCCS.

### Supplementary Tables

**Table S1. BTC expression classifiers**

| Reference | Classes | Classifier data source | DOI |
|---|---|---|---|
| Andersen *et al.* 2012 [7] | 2 | Supplementary Table 1 | 10.1053/j.gastro.2011.12.005 |
| Sia *et al.* 2013 [82] | 2 | Supplementary Table 2 | 10.1053/j.gastro.2013.01.001 |
| Ahn *et al.* 2019 [9] | 2 | Supplementary Table 1 | 10.1007/s12072-019-09954-3 |
| Montal *et al.* 2020 [83] | 4 | Table S11 | 10.1016/j.jhep.2020.03.008 |
| Nepal *et al.* 2021 [84] | 3 | Table S6 | 10.1016/j.jhep.2020.11.033 |
| Song G. *et al.* 2022 [10] | 2 | Supplementary Data 5 | 10.1038/s41467-022-29164-0 |
| O'Rourke *et al.* 2024 [8] | 2 | Table S2 | 10.1136/gutjnl-2023-330748 |
| Fan *et al.* 2024 [85] | 2 | Supplementary Figure 7 | 10.1038/s41467-024-44748-8 |
| Vijay *et al.* 2025 [11] | 4 | Table S5 \| RNA_cluster_signature | 10.1101/2024.07.04.601970 (bioRxiv) |

**Table S2. Top Scoring Pairs for expression consensus cancer subtypes (eCCS)**

| eCCS-A gene | eCCS-B gene |
|---|---|
| *DCDC2* | *KCNN4* |
| *CRMP1* | *CRYBG1* |
| *KLHL5* | *ACSL5* |
| *FXYD2* | *SLC44A4* |
| *KAAG1* | *TMPRSS4* |
| *BAAT* | *PRR15* |
| *BICC1* | *PTGS2* |
| *PDGFD* | *PLAC8* |
| *ANXA9* | *TSPAN1* |
| *RASSF8* | *MFSD2A* |
| *LDOC1* | *STYK1* |
| *HUNK* | *RHBDL2* |
| *MKKS* | *RPS6KA1* |
| *STRADB* | *TMEM164* |
| *ONECUT1* | *AGR2* |
| *EVC* | *PLAUR* |

**Table S3. Clinico-genomic characterization of samples classified as Other by the CCS framework.**

| Case ID | Presentation | Genomics | RNA PCA Outlier |
|---|---|---|---|
| BTC_0153 | SATB2+ (IHC) | *OCLN*::*RASGRF2* fusion | |
| BTC_0025 | | TMB > 100 | |
| BTC_0196 | large duct, mass forming iCCA | *CCNE1* rearrangement only | PC6 |
| BTC_0068 | HCC-CCA | *TP53*, *NTRK1* amplification | PC1, PC7 |
| BTC_0058 | Epithelioid malignancy, sarcomatoid | No drivers identified | many PCs |
| BTC_0124 | pCCA | HRD (*XRCC2* deletion) + *TP53* | PC5, PC6 |
| BTC_0056 | initial mass HCC-iCCA that underwent ablation and then recurred next to it as iCCA | *TP53* only | |
| BTC_0098 | | *CDKN2A* deleted only | |

**Table S4. Model weights for genetic consensus cancer subtype (gCCS), including chromosome arms as relative copy number compared to normal/control sample. VAF: Variant allele frequency.**

| Term | Coefficient |
|---|---|
| Intercept | -1.98 |
| 1q | 2.52 |
| 4p | 2.96 |
| 6q | -0.91 |
| 9q | -2.20 |
| 13q | -1.19 |
| 18q | 2.35 |
| 19p | 5.17 |
| Mean VAF | 1.92 |

**Table S5. Clinical characteristics of CCS-A and -B**

| | | Overall | CCS-A | CCS-B |
|---|---|---|---|---|
| | | n = 136 | n = 65 | n = 71 |
| **Age at Diagnosis** | | 59.7 (12.2) | 61.2 (10.8) | 58.4 (13.3) |
| | Unknown | 2 | 1 | 1 |
| **Sex assigned at birth** | | | | |
| | Female | 52 (39%) | 28 (44%) | 24 (34%) |
| | Male | 83 (61%) | 36 (56%) | 47 (66%) |
| | Unknown | 1 | 1 | 0 |
| **Stage at Diagnosis** | | | | |
| | I & II | 44 (40%) | 28 (51%) | 16 (30%) |
| | III | 20 (18%) | 7 (13%) | 13 (24%) |
| | IV | 45 (41%) | 20 (36%) | 25 (46%) |
| | Unknown | 27 | 10 | 17 |
| **Nodes at Diagnosis** | | | | |
| | N0 | 64 (53%) | 34 (58%) | 30 (48%) |
| | N1 | 42 (35%) | 19 (32%) | 23 (37%) |
| | N2 | 2 (1.7%) | 0 (0%) | 2 (3.2%) |
| | NX | 13 (11%) | 6 (10%) | 7 (11%) |
| | Unknown | 15 | 6 | 9 |
| **Metastases at Diagnosis** | | | | |
| | M0 | 77 (65%) | 38 (64%) | 39 (65%) |
| | M1 | 42 (35%) | 21 (36%) | 21 (35%) |
| | Unknown | 17 | 6 | 11 |

**Table S6. Small substitution drivers**

| gene | chr. | n | Element (FDR) | dN/dS (q) | LOH (FDR) | LOH n |
|---|---|---|---|---|---|---|
| *TP53* | 17 | 62 | 3.50E-72 | 0 | 1.30E-14 | 53 |
| *ARID1A* | 1 | 33 | 2.40E-09 | 0 | 0.099 | 24 |
| *BAP1* | 3 | 31 | 1.60E-27 | 0 | 2.80E-05 | 29 |
| *SMAD4* | 18 | 24 | 7.50E-16 | 0 | 2.60E-06 | 19 |
| *KRAS* | 12 | 21 | 2.40E-26 | 0 | 1 | 2 |
| *IDH1* | 2 | 19 | 1.10E-14 | 2.60E-11 | 0.72 | 0 |
| *EPHA2* | 1 | 19 | 2.40E-09 | 2.40E-06 | 0.5 | 14 |
| *PBRM1* | 3 | 17 | 3.40E-05 | 2.60E-08 | 0.025 | 15 |
| *STK11* | 19 | 12 | 6.10E-06 | 5.40E-06 | 0.0015 | 8 |
| *ELF3* | 1 | 11 | 0.0063 | 4.60E-07 | 1 | 1 |
| *TGFBR2* | 3 | 9 | 0.45 | 0.0034 | 0.0022 | 11 |
| *CTNND1* | 11 | 8 | 1 | 0.011 | 0.72 | 0 |
| *NRAS* | 1 | 7 | 5.30E-05 | 0.0036 | 0.5 | 0 |
